# Condensin-driven loop extrusion on supercoiled DNA

**DOI:** 10.1101/2021.05.15.444164

**Authors:** Eugene Kim, Alejandro Martin Gonzalez, Biswajit Pradhan, Jaco van der Torre, Cees Dekker

## Abstract

Condensin, a structural maintenance of chromosomes (SMC) complex, has been shown to be a molecular motor protein that organizes chromosomes by extruding loops of DNA. In cells, such loop extrusion is challenged by many potential conflicts, e.g., the torsional stresses that are generated by other DNA-processing enzymes. It has so far remained unclear how DNA supercoiling affects loop extrusion. Here, we use time-lapse single-molecule imaging to study condensin-driven DNA loop extrusion on supercoiled DNA. We find that condensin binding and DNA looping is stimulated by positive supercoiled DNA where it preferentially binds near the tips of supercoiled plectonemes. Upon loop extrusion, condensin collects all nearby plectonemes into a single supercoiled loop that is highly stable. Atomic force microscopy imaging shows that condensin generates supercoils in the presence of ATP. Our findings provide insight into the topology-regulated loading and formation of supercoiled loops by SMC complexes and clarify the interplay of loop extrusion and supercoiling.

## Introduction

Structural maintenance of chromosomes (SMC) complexes such as condensin and cohesin are key players for DNA organization in all organisms^1^. These SMCs organize chromosomes into loops as a result of DNA loop extrusion, a hypothesis that was directly confirmed by recent single-molecule experiments^2–6^. However, these *in vitro* studies typically used a straight piece of naked DNA, while DNA in cells is supercoiled, entangled, and bound by a myriad of DNA-binding proteins. Torsional stresses, which are inevitably induced by transcription and replication fork progression, cause DNA to adopt superhelical structures with locally under- or overwound DNA, known as negative or positive supercoiling, respectively. Such supercoiling typically involves large structural rearrangements of the DNA through the formation of plectonemes where the DNA helix is coiled onto itself (Fig.1a). It remains unknown how the loop-extrusion activity of SMC proteins deals with the supercoiling states of DNA that are abundant in cells.

Previous reports pointed to various DNA-supercoiling-specific interactions of SMC complexes. In yeast, mitotic DNA was found to manifest positive supercoiling that was dependent on the SMC2 subunit of condensin^7^, condensin was suggested to reduce transcription-induced underwound DNA^8^, and Chip-seq data suggested that cohesin can trap torsional stresses ahead of the replication fork^9^. In mammalian cells, it was shown that cohesin-mediated DNA loops induce topological stresses^10^. Gel-based^11,12^ and electron-imaging^13^ studies of condensin I from Xenopus egg extract showed an ATP-dependent generation of supercoiling in plasmids, and human condensin I showed similar results^14^. Other *in vitro* studies showed that yeast condensin^15^ and smc5/6^16^ can stabilize plectonemes on supercoiled DNA, while the cohesin heterodimer smc1/3 was reported to compact DNA in a supercoiling-dependent fashion^17^. Despite all these data, however, mechanistic insights are largely lacking and it remains unclear how these findings are linked to the recently found DNA-loop-extrusion activity of SMC proteins.

Here, we use real-time single-molecule imaging to study the dynamic interplay of DNA supercoiling and DNA loop extrusion by yeast condensin. We find that DNA binding and loop extrusion by condensin are stimulated by positive supercoiling. Condensin is found to preferentially bind to the tip of a plectonemes, whereupon it starts loop extrusion. Surprisingly, we find that loop extrusion by condensin absorbs all plectonemes on supercoiled DNA and locks them into a single supercoiled loop that is highly stable. The resulting supercoiled DNA loop subsequently acts as a favorable substrate for recruiting additional condensins. AFM imaging shows that condensin generates DNA supercoiling in an ATP-dependent manner. Finally, we discuss how our *in vitro* observations of the strong impact of DNA supercoiling on loop extrusion mechanics may impact DNA loop extrusion in cells.

## Results

### Condensin-mediated DNA loop extrusion removes nearby supercoiled plectonemes and generates one supercoiled DNA loop

To study the interaction between condensin and supercoiled DNA, we generated positively supercoiled DNA by adding intercalating Sytox orange (SxO) dyes onto torsionally constrained 42-kilobasepair (kbp) DNA substrates that were attached to a passivated surface at both ends with multiple binding groups^18,19^ (Fig. 1a). Visualization was done in HILO (highly inclined and laminated optical) microscopy. Similarly, negatively supercoiled DNA could be made by attaching pre-SxO-loaded DNA to the surface and subsequently reducing the SxO concentration^18^. Experiments were done on positively supercoiled DNA unless denoted otherwise.

**Fig. 1.**
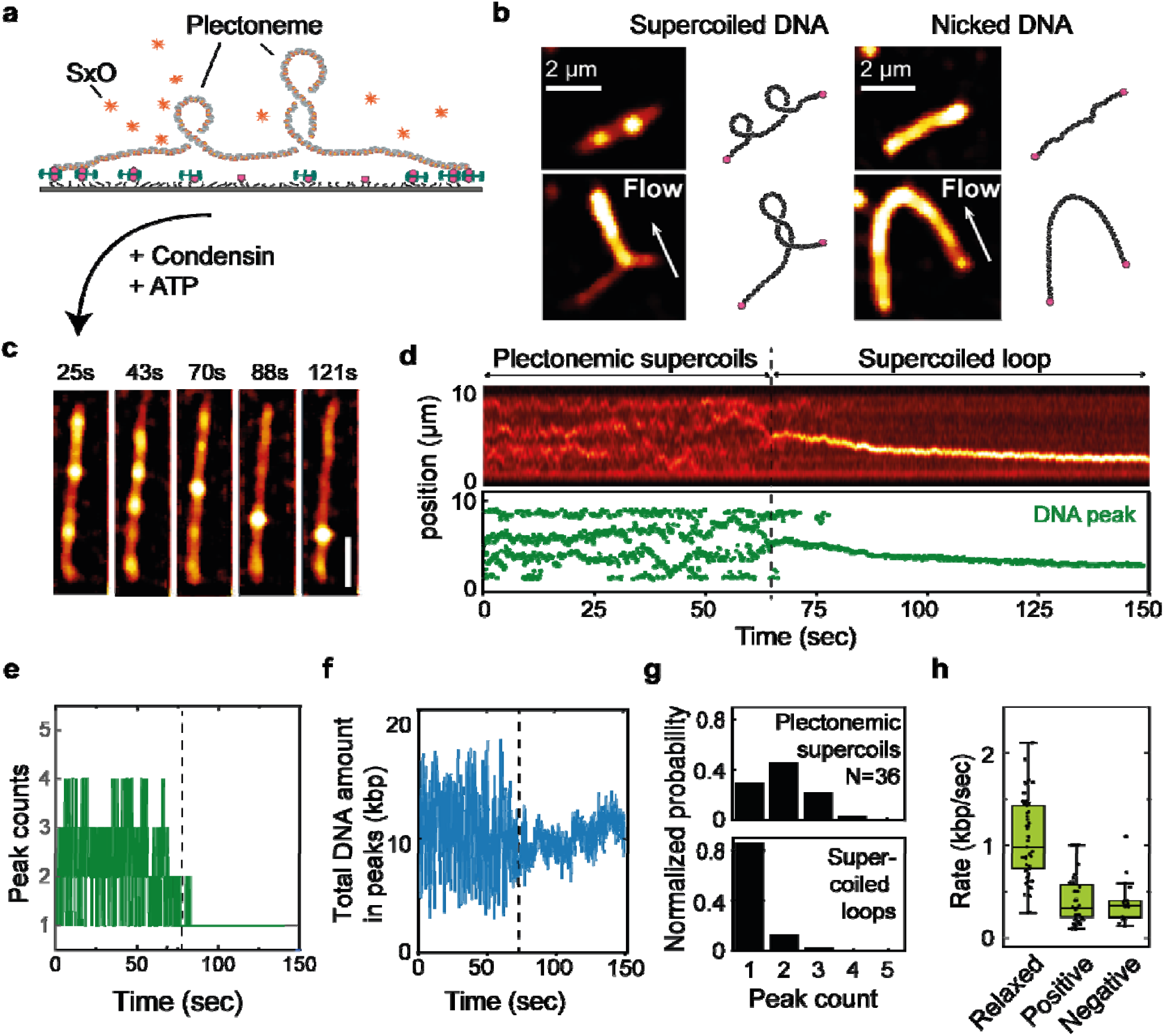
Condensin-mediated loop extrusion collects and stabilizes plectonemes into a single supercoiled DNA loop. (a) Schematic of intercalation-induced DNA supercoiling assay used for this study. (b) Snapshots showing single-molecule visualizations of double-tethered SxO-stained supercoiled DNA (left) containing multiple plectonemes (top-left) which can be stretched into a single plectoneme (top-bottom), and the images of the same nonsupercoiled molecule after nicking (right). Arrows indicate the direction of buffer flow. (c) Snapshots and (d) fluorescence-intensity kymograph (top) and the corresponding individual DNA peak positions (bottom), showing a loop extrusion event on supercoiled DNA. Magenta points indicate the position of DNA intensity peak that signals where loop extrusion occurs. (e) Corresponding number of total DNA peaks in (d). (f) The total amount of DNA within the DNA peaks in panel (c-d) verse time. (g) Comparison of the number of DNA peaks before (left) and after the loop extrusion (right) from N=36 molecules. (h) Rate of loop extrusion for relaxed DNA, positively, and negatively supercoiled DNA, from N=49, 36, 17 molecules, respectively.

Individual plectonemes were directly observable as local intensity maxima along the DNA, and their position, intensity, and number dynamically changed over time very frequently, consistent with previous reports^18–20^. Stretching these DNA spots sideways by applying buffer flow perpendicular to the attached DNA showed elongated DNA structures with a higher intensity than the rest of the DNA tether, indicating a plectoneme with two DNA helices that were mutually intertwined (Fig. 1b). Unlike the open loops that we observed for nicked DNA,^2^ we did not observe any temporary separation of the two arms, consistent with the plectonemic DNA structure. Upon nicking the same molecule with a prolonged exposure of excitation light, the plectonemes abruptly disappeared, the DNA image changed to exhibit a homogenous intensity, and side flow stretched the DNA into an arc shape – all confirming that the DNA relaxed to a torsionally unconstrained state upon nicking.

Upon addition of condensin (1-2 nM) in the presence of ATP (5 mM) to supercoiled DNA, the dynamic plectonemic state changed into a single spot that grew in intensity over time, see Fig. 1c,d. More specifically, at time t ~ 67 s in Fig. 1d, a single DNA intensity peak (magenta) appeared among the multiple plectoneme peaks, which gradually increased its size as it moved along the DNA, until it stalled (t ~ 120s). This observation is clearly reminiscent of the characteristic dynamics of condensin-mediated loop extrusion. Strikingly however, for this supercoiled DNA, all the plectonemes that were continuously present at t < 67 s disappeared upon the onset of the condensin-mediated DNA loop extrusion, whereupon all DNA intensity was concentrated at the loop location (t ~77s; Fig. 1d; Fig. s1 and video 1 for more examples). Figure 1e quantifies our observation and shows that 1-4 plectonemes (average 2.6) were present before t=67 s, whereas a single peak was observed after the loop extrusion (t ~77s). The final amount of DNA that was collected to the loop location was not much larger than the amount of DNA within plectonemes that were initially present (see Fig. 1f), indicating that condensin did gather the plectonemic DNA during loop extrusion but not substantially more. These data indicate that upon formation of a loop, condensin collects all the plectonemes along the 42 kilo-base pair DNA and stabilizes them at its location.

The observed behavior, i.e., that condensin-driven DNA loop extrusion leads to an abrupt transition from a dynamic plectonemic state to a stable loop by collection of all plectonemes into a single position, was consistently observed in nearly all loop extrusion events that we analyzed (N=32 out of 36, Fig. 1g), with a few exceptions where one or more plectonemes remained after loop formation (Fig. s2). Note that in the latter cases, condensin was bound near the tethered end of DNA and the (asymmetric) DNA loop extrusion was limited by reaching the end position of the DNA. Given that the linking number (a measure of the degree of supercoiling writhe) is conserved in a topologically constrained DNA molecule, the vanishing of all plectonemes along the two arms of the DNA molecule indicates that the local region of the extruded loop must be highly supercoiled.

We compared the speed of DNA loop extrusion between supercoiled and relaxed DNA (Fig. 1h), as obtained from the slope of the initial linear part of the loop growth versus time, cf. Fig.1d between 69 and 89 s. The observed average rate for supercoiled DNA (~0.4 kbp/s; both for positive and negative supercoiling) was almost a factor 3 lower than that of torsionally relaxed DNA (~1.1kbp/s). This may indicate that the resulting supercoiled topology of the DNA loop that is being extruded slows down the speed of loop extrusion.

### Condensin loads near a plectoneme tip and moves downward along the plectoneme upon loop extrusion

Where does condensin bind to supercoiled DNA and how does it relocate upon loop extrusion? To address these questions, we co-imaged DNA with condensin complexes that were labelled with a single fluorophore (ATTO647N). Interestingly, we observed that condensin often bound at a plectoneme, then diffused together with it along the DNA, whereupon it unbound without extruding a DNA loop (Fig. 2a, b). We note that such a temporary binding of condensin, which did not induce loop extrusion, was also reported for nonsupercoiled DNA^2^. The co-localization probability of condensin and plectoneme, estimated by counting the events where the positions overlapped within our diffraction limit (Fig. 2b; see Methods), revealed that at least 70% of all DNA binding events of condensins co-localized with plectonemes (N = 49; Fig 2c). This shows that condensin has a higher binding affinity to plectonemic supercoiled DNA compared to regular non-plectonemic DNA.

**Fig. 2.**
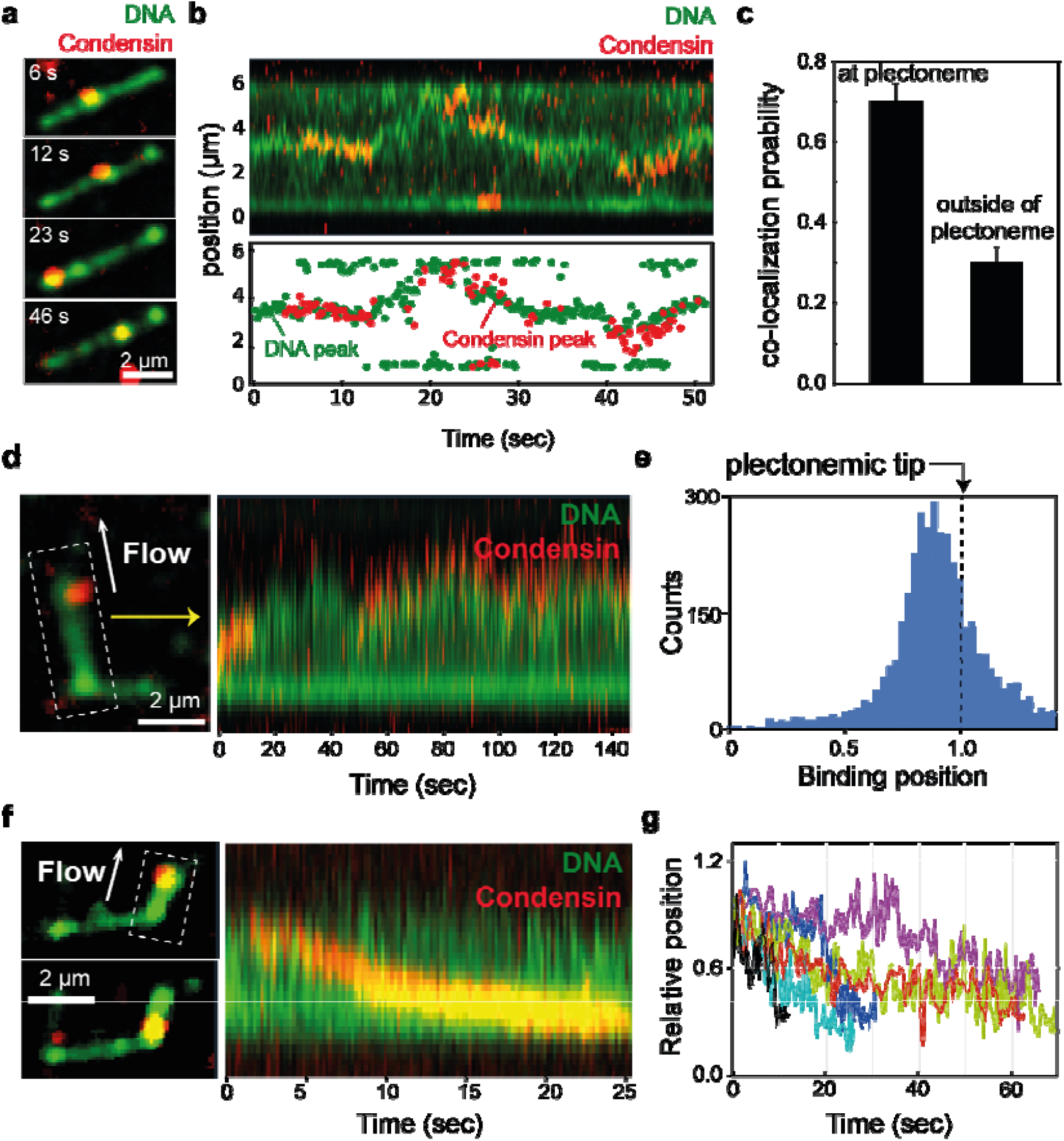
Condensin loads near a plectoneme tip, and moves downward along the plectoneme upon loop extrusion. (a) Snapshots and (b) a kymograph (top) and the associated positions of SxO-stained DNA and Atto647N-labeled condensin peaks (bottom), showing co-localizations of condensins and DNA plectonemes. (c) Co-localization probability of condensin and DNA plectonemes. Data shows the mean ± 95% confidence interval (N=49 molecules). (d) Snapshot (left) and kymograph (right) showing condensin localization near the plectoneme tip. Arrow indicates a direction of buffer flow. The dashed box indicates the area of the image included to build the kymograph. (e) Binding distribution of condensin along the length of DNA plectonemes estimated from the kymographs as in panel (d) for N= 27 molecules, where the scale 0-1 denotes the position from the stem to the tip along the plectoneme. (f) Snapshot (left) and kymograph (right) showing condensin moving from the plectoneme tip towards the stem. (g) Change in position for condensins that bound at the tip and moved along the length of DNA plectonemes extracted from the kymographs as in panel (f) for N= 6 molecules.

Subsequently, we measured where within a plectoneme condensin preferentially did bind, at its stem, middle, or tip. To this end, we quantified the binding locations of condensin complexes on side-flow-stretched plectonemes (Fig. 2d). Kymographs plotted along the length of a stretched DNA plectoneme clearly showed that the majority of condensins initially bound near the plectoneme tip (Fig. 2d). Quantification of the binding events (N=27 molecules; Fig.2e) showed a peak at a position of ~0.9, where a value 0 corresponds to the plectoneme stem and 1 to the tip location, indicating that the majority of condensins did bind in the vicinity of the plectoneme tip. This suggests that condensin favors binding at the apical loops of DNA plectoneme tip, instead of at DNA crossings at the plectoneme body or at the stem. Upon loop extrusion, the plectoneme-tip-bound condensins moved downwards and reached to the body or the stem of the plectoneme (Fig. 2f), as manifested by the changes of the condensin positions within the flow-directed plectonemes (Fig. 2i; N=6).

Next, we investigated where a condensin ends up on the plectoneme after loop extrusion, which we visualized by side flow after loop extrusion, see Fig. 3. In accordance to the observations of Figs. 1 and 2, condensin mostly bound to the tip of a plectoneme (N=25 out of 30 molecules; e.g. t =16 s in Fig. 3a, b; video 2) and gradually reeled in DNA while simultaneously localizing all the plectonemes near the condensin location (until t ~ 40 s; Fig. 3a, b). After the loop was formed, we applied a buffer flow (t=55 s) and monitored the location of the condensin along the supercoiled DNA loop (Fig. 3c, Fig. s3 for more examples). This revealed that, remarkably, condensin now was localized at the stem of the plectonemic loop, implying that plectonemic DNA was entirely absorbed into the supercoiled DNA loop by the condensin in the process of loop extrusion. Alternatively, we also observed condensins which extruded a supercoiled loop with similar dynamics (Fig. 3d, e), that ended up at a central location along the plectoneme (Fig. 3f, Fig. s4 for more examples). Even in these cases, all plectonemes in the DNA still were absorbed into the position of the extruded loop (t > 45 s; Fig. 3e). Localization of condensin occurred with similar frequencies at the stem or in the central body of the supercoiled loop after extrusion, while condensins rarely were found to end up located near the tip after loop extrusion (Fig. 3g). These data suggest that the condensin-mediated supercoiled loop does adopt plectonemes from outside the loop region to incorporate them inside the extruded loop or to stabilize them to a position just below the condensin complex.

**Fig. 3.**
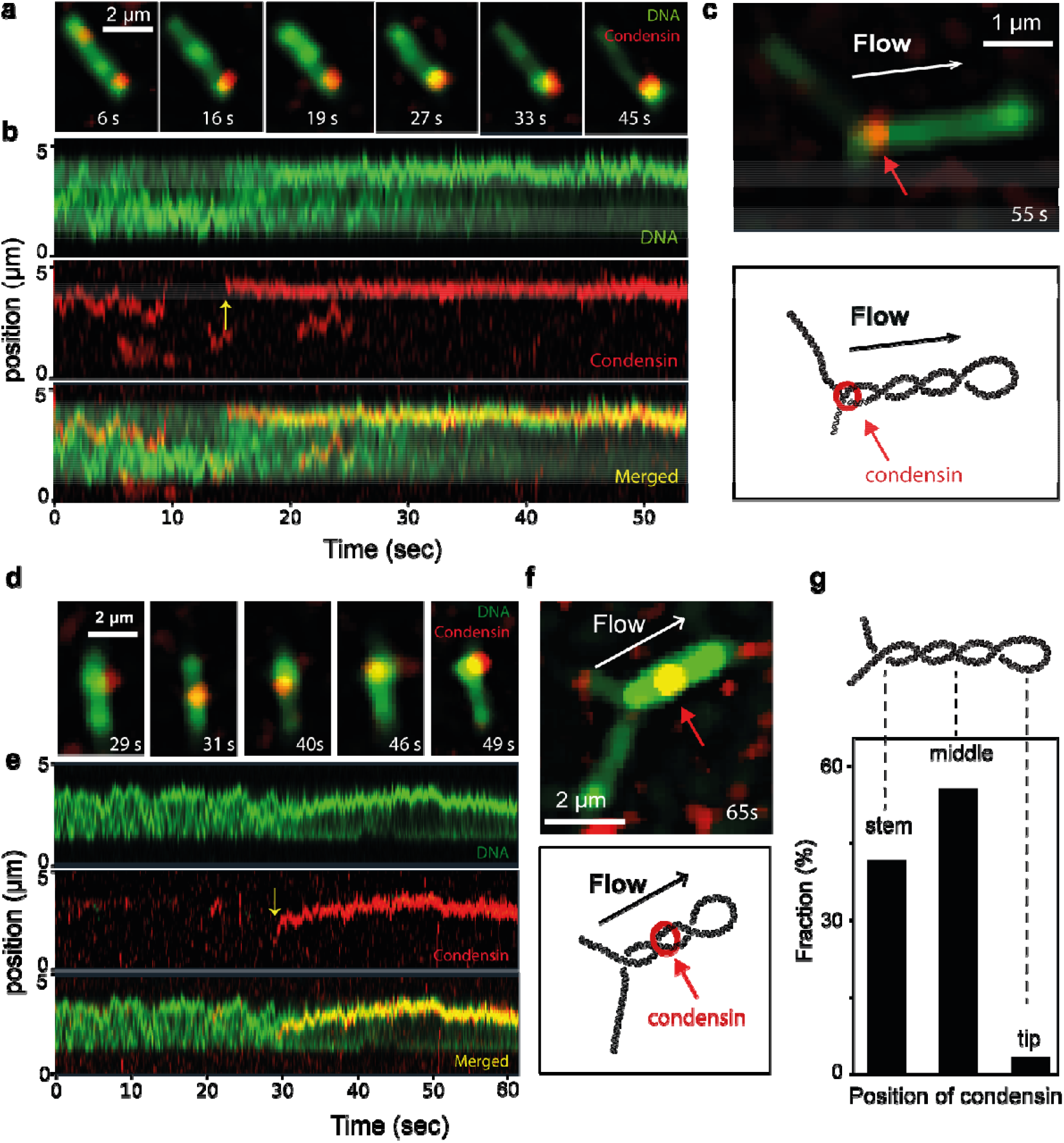
Condensin is located at the stem or at the middle of a plectoneme after extrusion of a supercoiled loop. (a) Snapshots and (b) kymographs showing loop extrusion on supercoiled DNA. Yellow arrow in (b) denotes the binding of condensin which subsequently led to loop extrusion. (c) Snapshot (top) and schematic (bottom) of a molecule in panel (a,b) in side flow, revealing the location of condensin along the plectoneme at the stem after extrusion of the supercoiled loop. (d) Snapshots and (e) kymographs and (f) the resulting snapshot with side flow of the supercoiled loop extrusion event, which shows a condensin located at the middle of a plectoneme after extrusion. (g) Fractions of occurrence of different condensin positions after extrusion of supercoiled DNA (N=36 molecules).

### Positive supercoiling promotes condensin loading and loop formation, while negative supercoiling does not

We further investigated whether the observed preferential loading of condensin on plectonemic DNA was dependent on the handedness of the DNA supercoiling. Interestingly, the probability to bind condensin to positively supercoiled DNA was found to be about 4-fold higher than to relaxed DNA (Fig. 4a). Remarkably, the binding affinity of condensin to negatively supercoiled DNA was similar to that to relaxed DNA (Fig. 4a), which indicates that condensin clearly favors binding to an overwound plectonemic DNA topology (positive supercoiling).

**Fig. 4.**
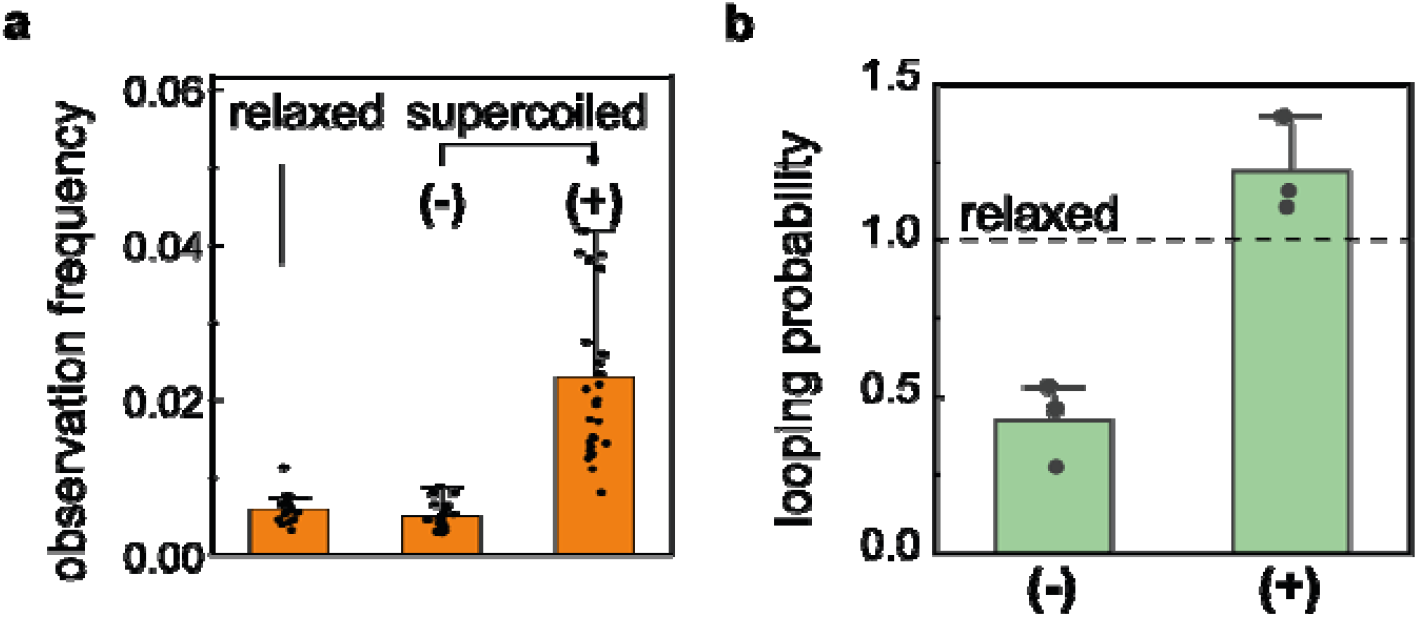
Positive supercoiling promotes condensin loading and loop formation, while negative supercoiling does not. (a) Probability to observe a condensin holocomplex on relaxed (nicked), negatively supercoiled, and positively supercoiled DNA. Number are calibrated as observation frequency per 1 kbp of DNA length (N=17, 26, 31 for relaxed, (−) and (+) coiled substrates, respectively). (b) Probability to observe loops on (−) and (+) coiled substrate relative to relaxed (nicked) DNA, mutually compared under the same conditions in the same field of view.

We then questioned whether the strong dependence of condensin’s binding affinity on the chirality of supercoiling also would affect the consequent loop formation probability. For this, we measured the fraction of all double-tethered DNA molecules that displayed a loop, and we compared the results for positively coiled DNA relative to that for relaxed DNA in the same field of view, and similarly for negative DNA supercoils compared to relaxed DNA. Interestingly, the estimated looping probability for negatively supercoiled DNA was significantly lower (~0.32) as compared to relaxed DNA, whereas for positive supercoils it was higher (~1.2; Fig. 4b). In other words, while positive supercoiling is promoting DNA loop extrusion, negative supercoiling is hindering it.

### Condensin-driven loop extrusion generates DNA supercoiling

To visualize condensin-mediated supercoiled loop with higher resolution, we used atomic force microscopy (AFM) and imaged the reaction products of condensin and topologically constrained DNA plasmids (2.96 kb in length) incubated with 1 mM of ATP, after drying the solution onto a mica substrate (Methods). The plasmids were first nicked and re-ligated with topoisomerase in order to make them torsionally constrained circular dsDNA with zero additional linking number with respect to the relaxed (nicked) form. Indeed, as Fig.5a,b shows, this led to a majority of molecules (60%) that showed a circular DNA without any crossovers or other signatures of supercoiling. Single crossovers (i.e., where a figure-0 plasmid was changed into a figure-8 shape) were observed also for these non-supercoiled plasmids, which can be attributed to the deposition of the DNA onto the mica surface in AFM sample preparation^21^. Adding the condensin SMC complex in the absence of ATP did not lead to an increase in the number of crossovers in the plasmids.

When adding condensin with ATP, however, a dramatic change was observed and the large majority of plasmids (72%; N=128) did get supercoiled, showing one or multiple crossovers (Fig.5a,b). The majority of plasmids now consistently exhibited plectonemic conformations that appeared as high-intensity DNA lines consistent with two dsDNA molecules that are tightly wound around each other. Quantification of the images (Fig. 5b,c) revealed that the initial plasmids showed only a ~30% fraction of entangled molecules (i.e. with crossovers or plectonemic conformations), whereas adding SMC and ATP strongly shifted this distribution to a much larger fraction of >70%. The data show that condensin can introduce a high degree of supercoiling into the plasmid DNA.

**Figure 5.**
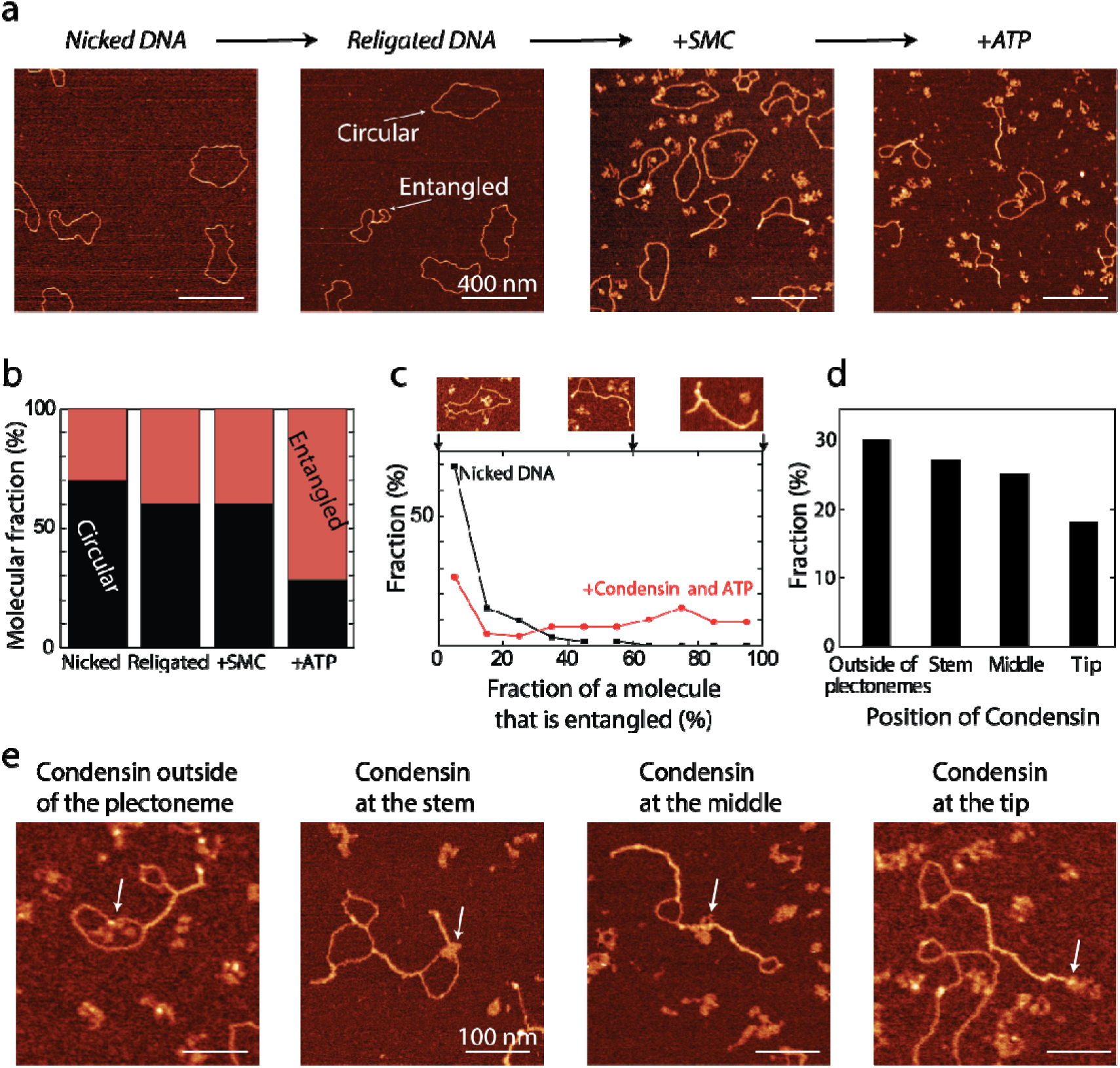
DNA supercoiling generation by condensin and ATP. a) Relaxed circular DNA molecules were nicked and religated, and incubated with condensin and ATP. The presence of condensin and ATP increased the number of DNA molecules that exhibit supercoiling. b) Population graph of circular and entangled molecules for various conditions (N=102, 128, 58, 92 molecules, respectively). c) Degree of entanglement for the nicked and +condensin+ATP samples. Images on top show examples of DNA that was entangled to different degrees. d) Binding position of condensin bound to supercoiled DNA molecules. e) Examples of condensin bound to supercoiled DNA molecules. White arrows denote the position of the condensin complexes.

Next, we quantified the position of condensins along the plasmids (Fig. 5d), where we classified their positions as outside the plectonemes, or at the tip, middle, or stem of plectonemes (see Fig.5e for examples). We found that, after addition of condensin and ATP, condensin distributed rather equally among all 4 categories, with a slightly smaller fraction for the location at the plectoneme tip.

Taken together, the AFM data show that the loop-extruding condensin complex generates DNA supercoils into plasmid DNA via an ATP-driven process. In other words, the data show that condensin does not only passively absorb and trap existing plectonemes along DNA, as already indicated by our fluorescence-imaging data (Fig. 3), but also that it can actively generate supercoils into DNA.

### Supercoiled DNA loops can recruit additional condensins

The preferential binding of condensin to DNA plectonemes (Fig. 2d) could lead to a situation where multiple condensins did bind to different plectonemes within the same DNA molecule. Interestingly, we observed that upon extrusion of a supercoiled loop, condensins that were bound at distant plectonemes could jointly be absorbed into one supercoiled loop. An example is presented in Fig. 6a, b where a condensin-driven supercoiled loop (t>16 s) did approach (t~20s) plectoneme-bound condensin(s) that were already present from t<16s, to eventually merge into a single location (t~30s). Subsequent imaging of the side-flow-stretched supercoiled loop after the merger (Fig. 6c; Fig. s3 for more examples) clearly showed multiple condensins that were bound along the supercoiled loop.

**Fig. 6.**
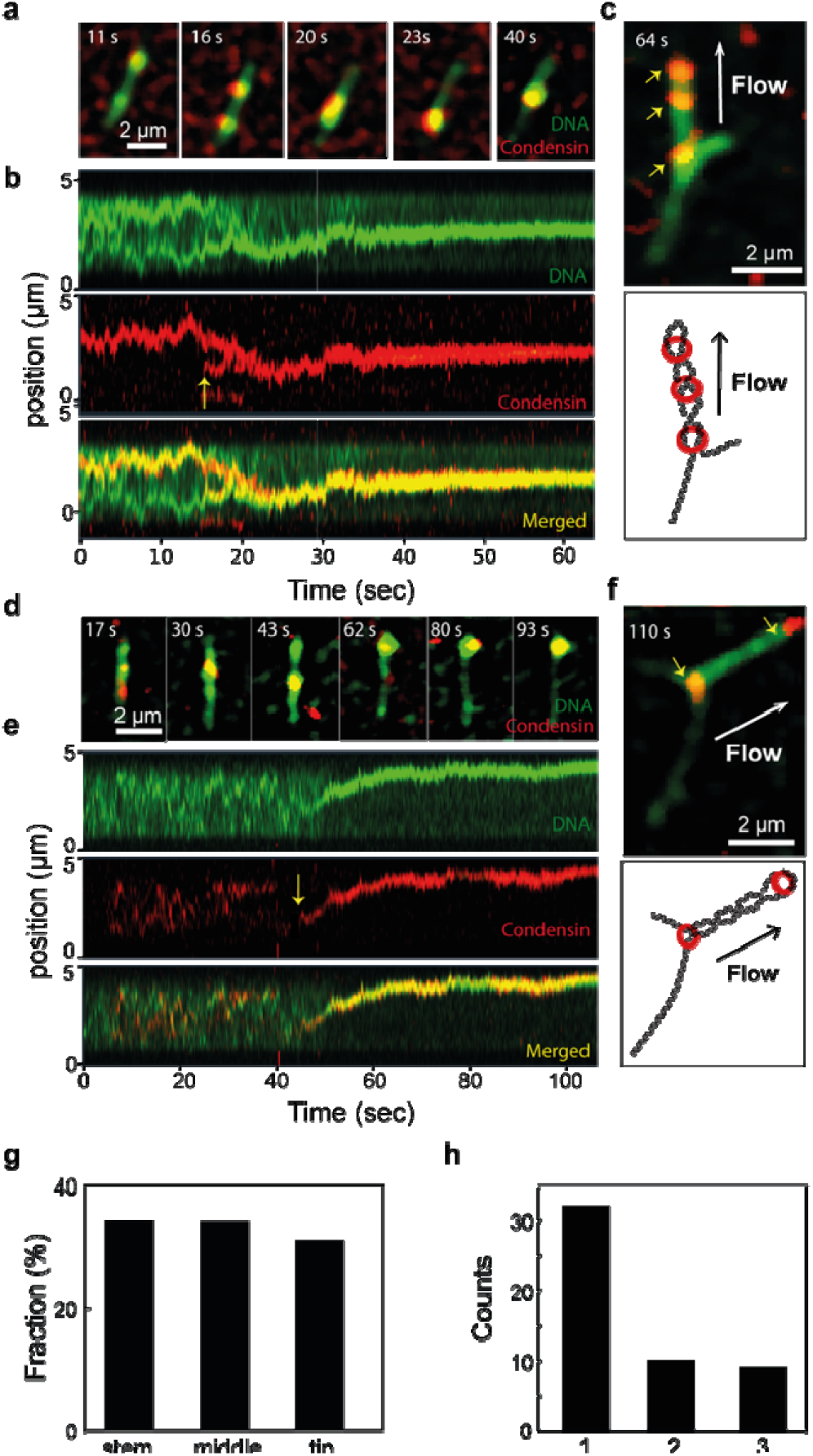
Recruitment of additional condensins onto a supercoiled loop. (a) Snapshots and (b) kymographs showing the merger of multiple condensins via supercoiled loop extrusion. Yellow arrow in (b) denotes the binding position of a condensin which subsequently lead to merging with another one(s). (c) Snapshot under side flow (top) and schematic (bottom) of the molecule in panel (a,b), revealing the location of multiple condensins along the plectoneme after the merger. (d) Snapshots and (e) kymographs and (f) the consequent snapshot under side flow of the supercoiled loop, which shows an additional condensin bound on the tip of an extruded supercoiled loop. Yellow arrows in (c, f) indicate the locations of condensins. (g) Occurrence of different positions of condensins along the plectomic loop after a supercoiled loop extrusion event for cases where we observed multiple condensins (N_tot_=61 for 26 DNA molecules). (h) Number of condensins on the supercoiled loop, estimated from photobleaching analyses.

**Fig. 7.**
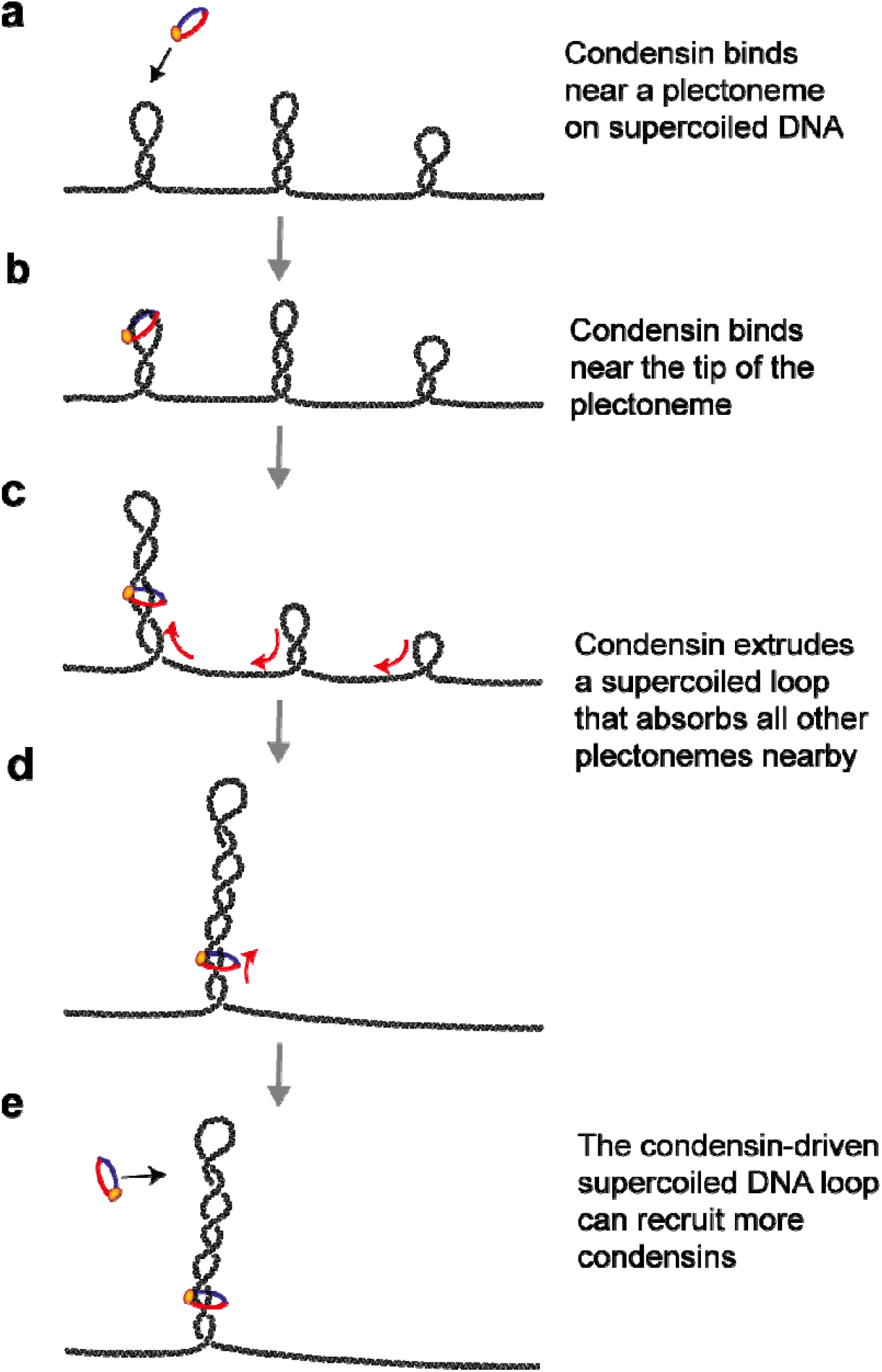
Model for condensin-mediated loop extrusion on supercoiled DNA. A condensin SMC complex loads at a DNA plectoneme (a), near its tip (b), and subsequently initiates the extrusion of a DNA loop that absorbs all neighboring plectonemes and locks them into a single supercoiled loop (c,d), which can be followed by the recruitment of additional condensins (e).

Furthermore, we found that a supercoiled loop acts as a favorable substrate for the recruitment of additional condensins (Fig. 6d, e). Consistent with the findings of Fig. 2d, these additional condensins often located near the tip of the supercoiled loops (Fig. 6f). Quantification of the locations of multiple condensins along supercoiled loops showed comparable numbers of condensins at the stem, in the middle, and near the tip (Fig. 6g), a trend consistent with the findings from AFM imaging (Fig. 5d). We estimated the number of condensin also by photo-bleaching experiments at the supercoiled loop location in the absence of buffer flow (Fig. s4). This showed that about 60% of all data (i.e., combined with data shown in Fig.3) involved a supercoiled DNA loop that contained a single condensin, while the rest were composed of 2 or 3 condensins on the loop (Fig. 6h).

## Discussion

In this study, we used time-lapse single-molecule imaging and AFM to study condensin-mediated loop extrusion on supercoiled DNA. Our findings provide a physical picture for the dynamic interplay of loop extrusion and DNA supercoiling (Fig.7), as follows. Condensin initially binds at the tip of a supercoiled plectoneme (a,b) and starts reeling-in the plectonemic DNA by loop extrusion. During loop growth, this emerging loop absorbs all the neighboring plectonemes and puts them into one supercoiled loop that is highly stable (c,d). The motor action of condensin will bring the complex downwards, where it ends up at a mid-position along the body or it reaches the stem, when eventually loop extrusion stalls due to the stalling tension within the DNA.^2^

Our single-molecule imaging showed that, surprisingly, the condensin-induced loop extrusion is clearly favoring positively supercoiled DNA over negatively coiled DNA. Condensin loads with a higher affinity onto positively supercoiled DNA, and it extrudes DNA loops more frequently. Our data are consistent with early gel-based findings that condensin binds more abundantly to positively coiled plasmids^12^. These *in vitro* results suggest that *in vivo* condensin recruitment may be stimulated in regions where positive supercoils are generated, e.g., ahead of transcription and replication machineries. This may explain the accumulation of condensins near transcription^8,22,23^ and replication sites^24^. Topology-enhanced loading likely is a more general phenomenon among SMC proteins. For example, cohesin and SMC5/6 have also been found to co-localize within regions where higher levels of catenation and torsional stress are present^9,25–27^

Our flow-induced visualization assay further showed that condensin initially bound preferentially near the plectoneme tip (Fig. 2d,e). We speculate that this may be due to the well-defined apical teardrop-shape loop structure at the tip of plectoneme which has a diameter of tens of nm^28^ that may serve as a favorable substrate for condensin, as it allows binding at two positions within the protein complex. Furthermore, the highly curved plectoneme tip may help condensin to start loop extrusion by circumventing the high energy cost associated with bending of DNA in the initiation of the loop extrusion.

Surprisingly, we found that the loop extrusion activity of condensin led to the absorption of all nearby plectonemes (Fig. 1 and Fig. 3). In other words, loop extrusion removes all supercoiling writhe from DNA and locks that into one localized spot – thus profoundly changing the DNA topology. This occurred over the full range of our 42 kbp DNA which provides a lower bound estimate for the range of the region where plectonemes are collected, since this length is limited in our assay. Our observations that loop extrusion incorporates existing intertwined structures does not support earlier ideas that an SMC loop extruder would act as a topological roadblock that would accumulate topological stresses ahead of the motor^25^, or that supercoiling itself can push SMCs^29^. The finding that loop extrusion localizes plectonemic supercoils in one defined location at the SMC may have interesting *in vivo* implications. It is tempting to speculate that the supercoiling domains found in cells^27,30,31^ may be organized by the plectoneme-localizing activity of SMCs. Interestingly, the average size of supercoiling domains (~100 kbp) found in human cells^30^ is of a similar order of magnitude as CTCF-defined loop domains (~180 kbp)^32^. Supercoiled chromatin loop domains may stimulate intramolecular interactions such as enhancer-promoter interactions^33^ or may act as efficient recruitment sites for topoisomerases by localizing topological stresses distributed along chromatin into defined regions. Indeed, various studies have reported a co-localization of topoisomerases and SMCs in both bacteria^34^ and eukaryotes^9,25^.

Our AFM data showed that yeast condensin with ATP can generate DNA supercoiling into plasmid DNA. This is consistent with previous *in vitro* gel-based and EM studies for condensin I complexes from frog^11,12^ and human^14^. Together with our findings from imaging experiments, this indicates that the supercoiled loop formation involves the active generation of supercoils in addition to the passive absorption of existing plectonemes. The DNA-supercoiling-generation activity of condensin raises the question on its implications for mitotic chromosome condensation. SMC-induced generation of DNA supercoils naturally condenses DNA as well as may facilitate decatenation of sister chromatids during anaphase onset^7,35^.

In summary, our study provides mechanistic insight into the dynamic interplay between condensin-mediated loop extrusion and plectonemic DNA supercoils. We observed a rich phenomenology, from topology-stimulated loading of condensin, absorption of plectonemic DNA via extrusion of supercoiled loop, to active generation of DNA supercoils. These findings provide novel insight into understanding loop extrusion of more complex DNA topologies, which is important as supercoiling is one of the fundamental properties of cellular DNA that helps constitute higher levels of chromosome organization.

## Acknowledgments

We thank E. van der Sluis and A. van den Berg for protein purification, and J. Kerssemakers, J.-K. Ryu, A. Katan, R. Barth, M. Tisma, Leon van Eendenburg for discussions.

## Author contributions

E. K. and C. D. designed the single-molecule imaging experiments, E. K. performed the imaging experiments and analyzed the imaging data, A. M. G. performed the AFM experiments and analyzed the AFM data. B. P. contributed in image analyses, J. van der T. prepared the DNA construct, C. D. supervised the work, all authors wrote the manuscript.

## Data and materials availability

Original imaging data and protein expression constructs are available upon reasonable request.

## Methods

### Condensin holocomplex purification and fluorescent labeling

We used our previously published expression and purification and labelling procotols^2,6^ to purify the pentameric *S. cerevisiae* condensin complex.

### Synthesis and purification of coilable 42kb DNA construct

A coilable 42kb DNA construct was made using linearized cosmid-I95^36^, containing on either end a Biotin-DNA-handle containing multiple biotins. The cosmid-I95 plasmid was amplified in Neb5 (New England Biolabs, C2987H) and the DNA was purified using a Qiafilter plasmid midi kit (Qiagen, 12243). Biotin-containing handles were made with a PCR on pBluescript SK+ (stratagene) with GoTaqG2 (Promega, M7845), in the presence of 1/5 biotin-16-dUTP (JenaBioscience, NU-803-BIO16-L) to dTTP (ThermoFisher, 10520651). The PCR was done with primer CD21 (GACCGAGATAGGGTTGAGTG) and CD22 (CAGGGTCGGAACAGGAGAGC), resulting in a 1238bp DNA fragment that contained multiple biotins. This was cleaned up using a PCR clean up kit (Promega, A9282). The Biotin-handle and Cosmid-I95 DNA were both digested for 2 hours at 37°C with SpeI-HF (NEB, R3133L) and subsequently heat inactivated for 20 minutes at 80°C. Resulting in linear ~42kb DNA and ~600bp Biotin-handles. The digested products where mixed together, where we use ~10 times molar access of the biotin-handle to linear cosmid-95. We then added T4 DNA ligase (NEB, M0202L) in the presence of 1mM ATP, overnight at 16°C and subsequently heat inactivated the next morning for 20minutes at 65°C. The resulting coilable 42kb DNA construct was cleaned up using an AKTA pure, with a homemade gel filtration column containing approximately 46ml of Sephacryl S-1000 SF gel filtration media (Cytiva), run with TE + 150mM NaCl_2_ buffer. The sample was run at 0.2ml/min and we collected 0.5ml fractions.

### Single-molecule visualization assay for studying condensin-mediated loop extrusion on supercoiled DNA

For immobilization of 42kb coilable DNA, we introduced 50 μL of ~ 1 pM of biotinylated-DNA molecules at a flow rate of 2 − 3 μL/min. Immediately after the flow, we further flowed 100 μL of a washing buffer (40 mM Tris−HCl, pH 8.0, 20 mM NaCl, 0.4 mM EDTA) at the same flow rate to ensure stretching and tethering of the other end of the DNA to the surface. We typically obtained a stretch of around 20−40% of the contour length of DNA.

To induce positive supercoiling of the tethered DNA, we flowed in 250 nM Sytox orange in condensin buffer (40 mM TRIS-HCl pH 7.5, 50 mM NaCl, 2.5 mM MgCl_2_, 1 mM DTT, 5% (w/v) D-dextrose, 2 mM Trolox, 40 µg/mL glucose oxidase, 17 µg/mL catalase) with 5 mM ATP. To prepare negatively supercoiled DNA, we first immobilized DNA to the surface in the presence of a high concentration of SxO (1 µM in condensin buffer with 5 mM ATP), and subsequently reduced the dye concentration to 250 nM for the measurements. The subsequent release of pre-bound SxO dyes after immobilization of the DNA results in negative supercoiling of the DNA.

Real-time observation of supercoiled-DNA loop extrusion by condensin was carried out by introducing condensin (1-2 nM) and ATP (5 mM) in the condensin buffer. Fluorescence imaging was achieved by using a home-built objective-TIRF microscopy. SxO-stained DNA and ATTO647N-labelled condensin were simultaneously imaged by alternating excitation of 532-nm and 640-nm lasers in Highly Inclined and Laminated Optical sheet (HILO) microscopy mode. All images were acquired with an EMCCD camera (Ixon 897, Andor) with a frame rate of 10 Hz.

### Data analysis for single molecule imaging

Fluorescence images were analysed using custom-written Python software. The noise from the images were removed with a machine-learning-based denoising method called “ Noise2void” as published before^37^. Fluorescence intensity kymographs were built from the intensity profiles from DNA and from condensin molecules per time point. Each vertical line on the kymograph (e.g. Fig. 1d) was obtained by summing fluorescence intensities of 11 pixels perpendicular to the DNA axis. Peaks on each vertical line from the kymograph were found with the peak-finding algorithm (“ scipy.signal.find_peaks”) in scipy^38^ which finds all the local maxima by comparing with the neighbouring values. The plectonemic peaks were further selected from the local maxima with 20 % threshold of the maximum peak prominences (relative peak intensities). This threshold removes the spurious low intense peaks.

Unlike non-loop extruding condensins that bind and unbind at a fixed position on DNA, condensins bound on plectonemic supercoiled DNA (e.g. Fig. 2a) diffuse along with the plectoneme^18,20^. In order to trace diffusing plectonemes and condensins, we consider peaks appearing in consecutive frames to be continuous if they appeared within 5 pixels (~375 nm or ~1 kbp) of each other. In order to reduce false positives due to noise, we only included peaks that were observed longer than for 10 consecutive frames. In this way, individual diffusing plectonemes and plectoneme-bound condensins could be tracked over time.

The estimation of the co-localization probability of condensin and DNA plectoneme peaks, shown in Fig. 2c, was performed as follows. From the tracked condensin and DNA peaks, we counted the peaks that had been in close proximity to each other (within 5 pixels, i.e. ~375 nm or ~1 kbp) for longer than 3 consecutive frames and divide this by the total number of tracked condensin peaks. For the data analysis regarding Fig. 3a, we obtained the probability to find a condensin peak per 1 kbp of DNA length by summing the total number of detected condensin peaks over all time points, and dividing this by the DNA length (42kbp) and by measurement time. For the analysis regarding Fig. 2e and 2g, we determined the position of condensin along the buffer-flow-stretched plectoneme by detecting the two ends of plectoneme (i.e. stem and tip of the plectoneme) per time point from the kymographs like Fig. 2d via fitting it with a super-Gaussian function. We then normalized the position of condensin peaks by the length of plectonemes at every time points. Note that since the condensin and DNA molecules were visualized at alternating time points that were 100 ms apart (alternative excitation mode), the normalized condensin position at every time point still can vary and result in a number larger than 1.

### Sample preparation and data analysis for AFM imaging

For the preparation of torsionally constraint DNA plasmid, pBlueScript SK+ (2961bp) DNA was purified from NEB5 at stationary phase with a qiafilter midiprep kit. To get constrained plasmid DNA without supercoils, we nicked the plasmid with nt.BspQI (NEB, R0644S), 2 hours 60°C and heat inactivated for 20 minutes 80°C. Some DNA was re-ligated overnight at 16°C in the same buffer with T4 DNA ligase, in the presence of 1mM ATP. Both the nicked and re-ligated plasmid DNA was subsequently purified using a PCR clean up kit.

Condensin stock solution was first diluted to a concentration of 25 nM. Then, the condensin was diluted into a volume of 20 µl containing 0.5 nM DNA, 1-2 nM Condensin in 50 mM Tris-HCl pH 7.5 and 50 nM NaCL. The condensin-DNA solution was incubated for 5 minutes. Afterwards, the solution was supplemented with MgCl_2_ to a final concentration of 5 mM and deposited onto a freshly cleaved mica. After 30 seconds, the surface was thoroughly washed with 3 ml of Milli-Q water and dried under nitrogen air-flow^39^.

Images were taken with a multimode-2 AFM from Bruker (Bruker corporation, Germany) using Scanasist-Air-HR tips from Bruker. The AFM was operated using Peak-Force tapping mode for imaging in air, at room conditions. Image data and processing (plane subtraction and flattening) was done with Gwyddion software.

